# β-bursting as a sensitive neural marker of inhibitory control in healthy older adults: a linear mixed-effects modelling and threshold-free cluster approach

**DOI:** 10.1101/2025.06.21.660835

**Authors:** Aliya C. M. Warden, Damian Cruse, Craig McAllister, Hayley J. MacDonald

**Affiliations:** School of Sport, Exercise and Rehabilitation Sciences, College of Life and Environmental Sciences, University of Birmingham, Birmingham, UK; Centre for Human Brain Health, University of Birmingham, Birmingham, UK; School of Psychology, College of Life and Environmental Sciences, University of Birmingham, Birmingham, UK; Department of Biological and Medical Psychology, University of Bergen, Bergen, Norway

**Author notes:** Corresponding author: Hayley J MacDonald, Address: Jonas Lies vei 91, 5009 Bergen, Norway.

**Keywords:** β-bursts, β-oscillations, electroencephalography, impulse control, response inhibition

## Abstract

Inhibitory control is essential for adaptive behaviour and declines with age, yet the underlying neural dynamics remain poorly understood. The β-rhythm (15–29 Hz) is thought to reflect inhibitory signalling within the fronto-basal ganglia network. Recent evidence suggests that transient β-bursts support inhibitory performance, often masked by conventional analyses of trial-averaged β-power. To reveal the link between trial-by-trial β-bursting and inhibition, we applied a recently developed analysis framework combining linear mixed-effects modelling (LMM) with threshold-free cluster enhancement (TFCE) during response inhibition and initiation in older adults. Twenty healthy older adults performed a bimanual anticipatory response inhibition task, while electroencephalography and electromyography were recorded to capture β-activity (β-burst rate/duration; averaged β-power) and muscle bursting dynamics, respectively. Our analysis revealed distinct β-bursting signatures absent in averaged β-power data. Following the stop-signal, parieto-occipital β-bursting presented before a temporal cascade from attentional to inhibitory processes. In addition to expected right fronto-central and bilateral sensorimotor activity, we observed left prefrontal β-bursting, indexing broader inhibitory network engagement during bimanual response inhibition. Moreover, we established a functional link between right sensorimotor β-bursting and muscle bursts during stopping, indicating rapid cortical suppression of initiated motor output. These results help clarify the mechanistic role of β-oscillations and underscore the sensitivity of β-bursting to both the timing and context of inhibitory demands in healthy older adults. Future research will help establish the potential of β-bursting, combined with LMM-TFCE analysis, as a clinically relevant marker of impulse control dysfunction.

**Significance statement:** Our novel application of an advanced statistical framework revealed distinct spatiotemporal β-bursting patterns during response inhibition and response withholding in healthy older adults, which were not captured by averaged β-power. Identifying a further link between cortical β-bursting and muscle-level suppression, the findings offer a mechanistic account of how the brain halts action in real time in older adults. This work provides a sensitive, trial-level framework for studying β-bursting measures in general, as well as inhibitory control across aging and clinical populations.

## Introduction

Supressing inappropriate actions is essential for goal-directed behaviour and widely assessed through response inhibition paradigms (Verbruggen et al., 2019). Such tasks recruit a right-lateralised fronto-basal ganglia network (Aron et al., 2016; Aron et al., 2014). Conventional Go/No-Go and Stop-Signal tasks probe withholding or cancellation of externally cued responses and reveal age-related slowing of inhibition (Rey-Mermet & Gade, 2018). However, these paradigms cannot capture the internally timed nature of everyday actions. The increasingly popular Anticipatory Response Inhibition Task (ARIT; He et al., 2022) instead requires withholding and cancellation of internally generated responses. Specifically, response withholding is proactively maintained and then strategically released for response initiation synchronised with a stationary target on Go trials, while Stop trials trigger reactive inhibition of the anticipated response via a stop-signal. Directly contrasting Go and Stop trials differentiates proactive versus reactive inhibitory control (Meyer & Bucci, 2016).

β-oscillations, a hallmark of motor-related neural activity, are positively correlated with GABAergic inhibition (Groth et al., 2021). Sensorimotor β-power decreases during movement (movement-related desynchronisation) and transiently increases afterwards (post-movement rebound) (Kilavik et al., 2013). However, raw β-band activity is better characterised by transient, burst-like events (<150 ms) rather than overall power (Sherman et al., 2016). In healthy young adults, sensorimotor β-bursting is thought to reflect an inhibited motor state, which must be released to initiate movement, but can be rapidly reinstated following frontal control signals during stopping. Response withholding is therefore characterised by inhibitory sensorimotor β-bursting, which progressively reduces for successful response initiation (Little et al., 2019; Wessel, 2020). Conversely, response inhibition features fronto-central β-bursting, which trigger inhibitory control processes through subsequent bilateral sensorimotor β-bursting (∼25 ms later) (Enz et al., 2021; Wessel, 2020). Increased fronto-central β-burst rate (Wessel, 2020) and volume (burst duration × frequency span × amplitude; Enz et al., 2021) predict successful stopping, while earlier single-trial β-bursts are associated with faster stopping (Hannah et al., 2020). Collectively, fronto-central and sensorimotor β-bursting patterns offer valuable insights into neural-behavioural associations underlying response withholding and inhibition.

Muscle-level measures can capture the downstream effects of these β-bursting neural dynamics. During the ARIT, healthy older adults display premature muscle bursts when withholding the planned response in ∼17% of Go trials and partial bursts on ∼50 % of successful Stop trials, reflecting initiation and rapid cancellation of a response before being executed (Warden et al., 2025). Partial burst latency provides a single-trial index of stopping (CancelTime; Raud et al., 2022). Right frontal β-bursting precedes CancelTime by ∼37-54 ms, with earlier β-bursts associated with faster CancelTime, establishing a preliminary link between frontal β-bursting and motor suppression (Jana et al., 2020). However, the mechanistic relationship between β-bursts across the wider inhibitory control network and these muscle responses at the single-trial level remains unclear.

Current analyses of β-bursting dynamics often rely on subject-wise averages of β-activity that may obscure meaningful trial-level and inter-individual differences - particularly in older populations with heterogeneous neural responses (Riccardi et al., 2025). The present study therefore adopted a recently developed analysis framework combining linear mixed-effects modelling (LMM) and threshold-free cluster enhancement (TFCE) (Visalli et al., 2024). LMM enables modelling of β-features at the single-trial level while accounting for within- and between-participant variability. Complementing this, TFCE enhances sensitivity to focal and distributed effects, enabling detection of subtle yet spatially extended β-activity. These techniques may therefore offer a more statistically robust approach for characterising β-burst dynamics, especially in older adults.

The first aim of this study was to apply combined LMM and TFCE (LMM-TFCE) techniques to contrast β-burst activity during response inhibition versus initiation in healthy older adults performing the ARIT. We hypothesised that increases in right fronto-central β-bursting would be followed by bilateral sensorimotor β-bursting during response inhibition. To investigate downstream effects of β-bursting, we applied our analysis framework towards a second aim of characterising the potential functional link between β-bursting and muscle bursting during response inhibition and withholding.

## Materials and Methods

### Participants

Twenty healthy older adults (69.3 ± 5.8 years, 9 female, 18 self-reported as right-handed) were recruited as part of a larger pre-registered study (see https://doi.org/10.17605/OSF.IO/KC2H3 and https://doi.org/10.17605/OSF.IO/W2FHP). Of note, the larger study includes comparisons between people with Parkinson’s disease and healthy older adults. Here, we have reported our novel application of advanced EEG/EMG analyses conducted on healthy older adults only.

Participants were recruited via the Birmingham 1000 Elders group and posters placed within local community hubs. Participants were recruited if aged 40-80, had no history of neurological conditions and had no reported vision impairment that was not corrected (e.g., with glasses). Participants were also required to score >23 on the Montreal Cognitive Assessment (MoCA; *M* = 27.4 ± 1.6) (Karlawish et al., 2013).

All participants gave written informed consent and received monetary compensation for their participation. The study received favourable opinion from an NHS Research Ethics Committee (South-East Scotland REC 02, IRAS ID: 328075) and was conducted in accordance with the Declaration of Helsinki.

### Behavioural task

As described previously (Warden et al., 2025), the current study employed a bimanual version of the ARIT (Fig. 1), programmed in MATLAB (R2019b, The MathWorks) and operated via custom microswitches. Participants were seated ∼0.6 m from a monitor displaying two vertical indicators, resting their forearms on a table mid-way between supination and pronation. The medial aspect of each index finger was used to lightly depress the two switches, with the left and right indicators corresponding to the respective fingers. After a 2 s delay, both indicators started to move upward from the bottom at equal rates.

**Figure 1.**
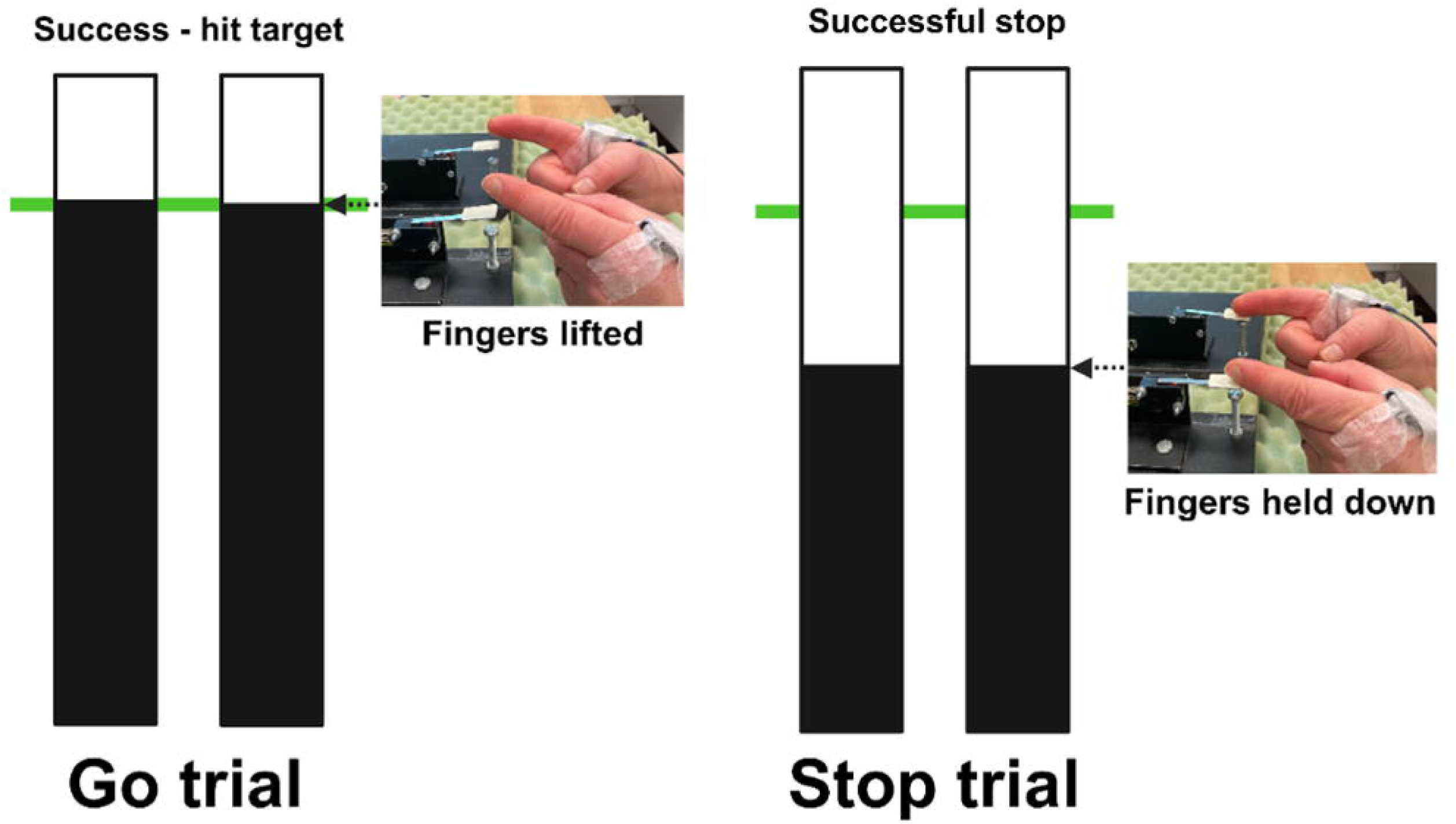
The Anticipatory Response Inhibition Task. During Go trials, participants initially held their index fingers on the respective switches and were instructed to lift their fingers once rising indicators reached the target line (at 800 ms). During Stop trials, the indicators stopped prematurely before the target and participants were instructed to inhibit the prepared finger lifts.

The task was primarily composed of Go trials (67%, 320 trials), where participants released both switches to intercept the rising indicators with a stationary target line at 800 ms (Fig. 1 left). Go trials were classed as ‘on-target’ if both indicators were within ±50 ms of the target line. During Stop trials (33%, 160 trials), both indicators stopped before reaching the target (Fig. 1 right), cueing participants to inhibit the prepared bimanual movement. As is common practice (He et al., 2022), an adaptive staircase algorithm was incorporated into Stop trials, dynamically changing the stop-signal delay (SSD) based on trial performance to keep task difficulty consistent across participants. The 2:1 trial ratio helped ensure that Go trials were the primed response. Immediate visual feedback reinforced performance, and participants were instructed to respond promptly and accurately to prevent strategic slowing, while maintaining relaxed muscles until response initiation to reduce baseline EMG noise.

### Recording and preprocessing

#### Electroencephalography

Continuous scalp-EEG was recorded with a 128-channel Biosemi ActiveTwo system (Biosemi, Amsterdam, The Netherlands). Two additional reference electrodes were recorded from the mastoid processes. EEG data was digitised at a sampling rate of 1024 Hz and preprocessed using a combination of custom scripts, the EEGLAB toolbox (Delorme and Makeig, 2004; http://sccn.ucsd.edu/eeglab) and the Fully Automated Statistical Thresholding for EEG artifact Rejection procedure (FASTER; Nolan et al., 2010; http://sourceforge.net/projects/faster). The data were first digitally filtered between 0.5 and 40 Hz, referenced to the average of the mastoids, and segmented into epochs from 500 ms before the bars started to rise until the end of the trial (+1000 ms).

Artefact rejection was performed using automated procedures based on FASTER (Nolan et al., 2010). First, channels with absolute z-scores >2.5 on any of the following measures were removed: voltage variance, mean correlation with other channels, and Hurst exponent. Second, trials containing non-stationary artifacts were excluded using the same threshold, based on mean voltage range across channels, mean voltage variance, and deviation of trial average voltage from the grand average. Third, independent component analysis was conducted on the remaining data (via the EEGLAB runica algorithm) and the ADJUST toolbox (Mognon et al., 2011) was used to automatically identify and remove components with the expected spatial and temporal features of blinks, eye-movements, and generic discontinuities. Upon visual inspection, any trials with artefacts that were not effectively cleaned by the above procedure were discarded. Fourth, any previously removed channels were interpolated back into the data. Finally, all data were re-referenced to the average of all channels. After data exclusions, a median of 159 on-target Go trials (range: 74-245) and 80 successful Stop trials (range: 65-83) were included in the analysis.

To enhance spatial resolution for β-burst detection, EEG data were transformed using the current source density method (CSD; Kayser and Tenke, 2006; https://psychophysiology.cpmc.columbia.edu/software/CSDtoolbox/index.html).

#### Electromyography

The lift response required participants to abduct their index fingers with the first-dorsal interosseous (FDI) muscle acting as a primary agonist. To measure this muscle activity, surface EMG was recorded via the Biosemi ActiveTwo amplifier (Biosemi, Amsterdam, The Netherlands) using two flat-type active electrodes placed over the FDI muscle of each hand, sampled at 1024 Hz. The scalp reference electrodes (CMS/DRL) were used as a ground.

The difference between the two electrode signals was amplified, bandpass filtered (50–450 Hz), and processed with custom MATLAB scripts (R2019b, The MathWorks) and EEGLAB. During preprocessing, EMG data were segmented into 1 s epochs from when the indicators started to rise until the end of each trial.

### Experimental design and statistical analysis

#### Task performance

Go trials without a lift response were excluded from all analyses. Lift-times on Go trials were recorded for each index finger via registration of the lifted switches. For each side, mean lift-time was calculated after trimming outliers (± 3 SD) (MacDonald et al., 2014). Stop-signal reaction time (SSRT) was calculated using the integration method, estimating the time needed to inhibit the response on Stop trials (Verbruggen et al., 2019). Trimmed lift-times averaged across side for Go trials were rank ordered and the nth lift-time selected, with n obtained by multiplying the number of lift-times by the probability of a response on a Stop trial. The time at which the staircase procedure stopped the indicators to achieve 50% success (*staircased* SSD) was subtracted from the nth lift-time.

### β-features

#### Time-frequency decomposition

Time-frequency decomposition was performed using the same general method as described by Enz et al. (2021) and Wessel (2020). Power estimates were computed across 15 linearly spaced frequencies from 15 to 29 Hz, using complex Morlet wavelets with 4 to 10 cycles logarithmically spaced. The squared magnitude of the convolved signal was used to obtain power for each electrode and epoch.

#### Average β-power

For wider comparison to earlier studies on β-oscillations during response inhibition, average β-power was calculated. Baseline activity was calculated as average β-power while the switches were pressed down, 400–100 ms before the indicators started to rise. Power estimates were then converted to decibels (dB) relative to this baseline period (baseline normalised).

#### β-burst detection

Transient β-bursts were identified from the raw (non-baseline normalised) power estimates via an established thresholding method (Enz et al., 2021; Hannah et al., 2020; Wessel, 2020). Peaks of β-activity exceeding 2 x median of the baseline (400–100 ms before the indicators started to rise) were detected as β-bursts, posited by Enz et al. (2021) as a sensitive threshold for investigation of the stopping process. The properties of identified β-bursts, including their incidence, timing and duration, were subsequently identified on a trial-by-trial basis during on-target Go and successful Stop trials.

### Spatiotemporal analysis

For successful Stop trials, an individualised time window (312.5–0 ms prior to SSRT) was extracted based on each participant’s SSRT. Trials with an SSD exceeding 700 ms were excluded to encompass the full range of SSRT values. This individualised time-locking aligns β-features more consistently across the study group, as they are temporally anchored to each participant’s stop-signal processing dynamics. For on-target Go trials, a window of equal length was taken relative to the earliest lift-time across hands on each trial, capturing β-activity as each participant prepares to respond.

To assess changes in β-activity during the response withholding and inhibition phases, 10 equal time bins of 31.25 ms length were defined (i.e., 32 samples at 1024Hz), with 0 ms representing the SSRT/lift-time. β-power was computed over whole epochs before being subdivided into time bins. β-burst rate was calculated as the number of β-bursts in each time window at each electrode. β-burst onset and offset were extracted at peak frequency and β-burst duration was calculated at this point (Hannah et al., 2020).

To visualise the topographical distribution of β-activity over the course of the trial, β-power, β-burst rate and β-burst duration were plotted across time bins for on-target Go versus successful Stop trials in a topographical grid representing the scalp surface (see Fig. 2).

**Figure 2.**
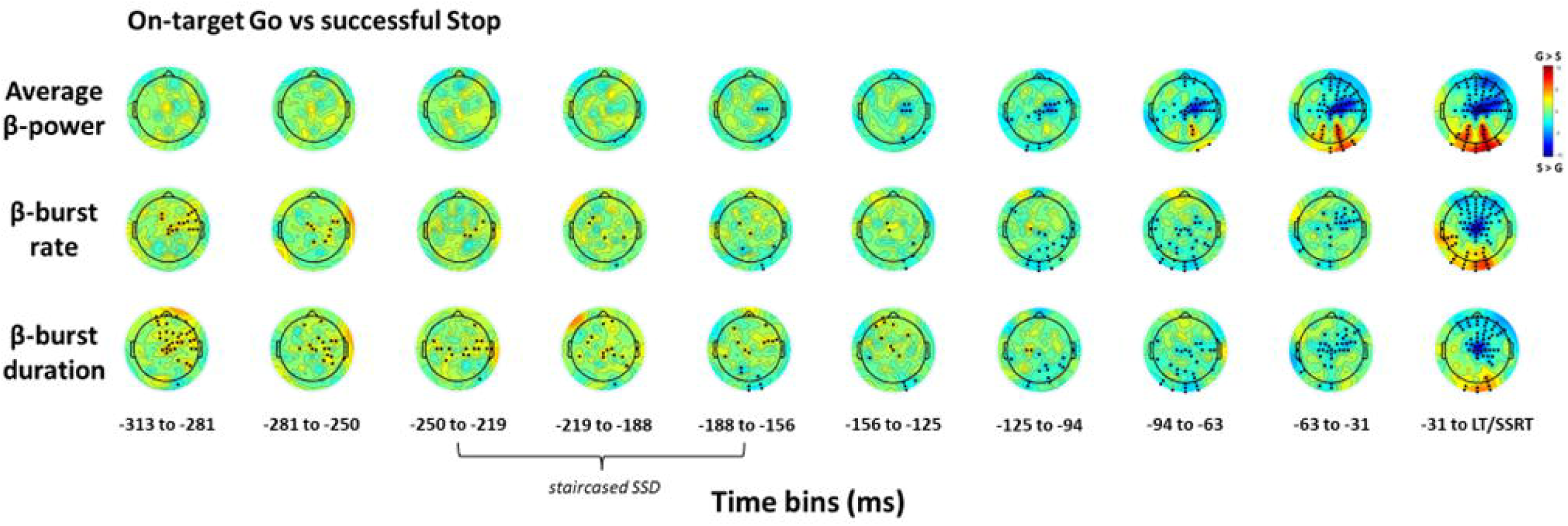
Topographical comparison of β-features between Go and Stop trials. Topoplots of T-statistics for β-power, β-burst rate and β-burst duration across ten ∼31 ms time bins leading up to the lift-time (LT) or stop-signal reaction time (SSRT) on on-target Go trials versus successful Stop trials. β-band frequency range was 15-29 Hz and β-bursts were detected using a 2-x median power threshold. As per the colour-bar, red indicates a relative increase in β-activity in Go trials (G) while blue indicates a relative increase in Stop trials (S). Black dots refer to significant electrodes following threshold-free cluster enhancement permutation testing (p < .05). SSRTs ranged from 180-239 ms, with staircased stop-signal delay (SSD) values falling across several timepoints, as indicated.

### Muscle bursting

EMG data processing was performed using MATLAB (2022a, The MathWorks). Trials with baseline root-mean-squared (rms) EMG exceeding 30 µV between 300–400 ms into the trial were excluded from the analysis. As per the second study aim, ineffective bursts of muscle activity (i.e. those that did not trigger a lift response) were identified on a trial-by-trial basis across Go and successful Stop trials.

Muscle bursts were defined as instances when the smoothed rectified EMG amplitude surpassed 15 SD of the baseline (rmsEMG at 0-400 ms) and remained above threshold for ≥5 ms. On Go trials, the main burst generating the lift response was identified as the last burst with an onset occurring before the recorded switch release. Accordingly, *premature bursts* were identified as suprathreshold EMG activity prior to the main lift response. On successful Stop trials, *partial bursts* represented muscle activity which decreased in amplitude before generating sufficient force to trigger the lift response. Bursts occurring within an individualised time window (±3 SD of trimmed Go lift-times) were classed as partial bursts, indicating preparation to respond. CancelTime was calculated as the difference in milliseconds between the SSD and peak amplitude of the partial burst (e.g., when muscle activation begins to decrease) (Raud et al., 2022).

The precise onset and offset times of these muscle bursts were automatically detected using a single-threshold algorithm (Hodges & Bui, 1996). Bursts separated by <15 ms were merged into a single burst. Detected bursts were manually reviewed on a trial-by-trial basis. Stimulus artifacts, and erroneous or unclear bursts (e.g., after lift-time, no clear separation with main burst) were de-selected at this stage and thus excluded from subsequent analyses. Burst trial incidence was calculated as % of traces with ≥1 burst.

### Linear mixed effects modelling

To examine β-activity during response inhibition versus initiation (aim 1), LMMs were developed using the lmeEEG toolbox (Visalli et al., 2024) in MATLAB (2024b, The MathWorks) for each β-feature (β-power, β-burst rate, β-burst duration). Briefly, this method fits a LMM to each electrode-by-timepoint, modelling and removing random effects (intercepts only), before conducting mass linear regressions on the marginal data. This approach allows for control of multiple comparisons by permutation testing without the enormous computational costs of fitting large numbers of LMMs. For each LMM, participant was included as a random effect and trial type (on-target Go versus successful Stop) was included as a fixed effect, with effects coding used for each contrast:

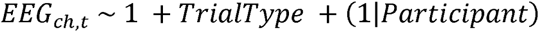

This statistical approach was utilised to effectively address both within- and between-participant variance, modelling β-activity on an individual trial basis, thereby accounting for its inherent variability. Controlling for random effects in this manner more accurately models the clustered system (Visalli et al., 2024), helping to isolate distinct β-activity patterns during response initiation versus inhibition processes.

To explore potential associations between β-bursting and muscle bursting (aim 2), two LMMs were developed on β-bursting dynamics (rate/duration) during Go/Stop trials which did or did not feature muscle bursting. Specifically, for Go and Stop trials separately and for burst rate and burst duration separately (as extracted above), we included the presence/absence of a muscle burst as a fixed effect and participant as a random effect:

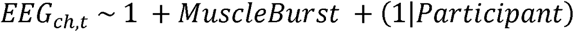

### Threshold Free Cluster Enhancement

Following linear mixed-effects modelling, TFCE (Mensen & Khatami, 2013) permutation testing was performed to control for multiple comparisons and identify significant (*p* < .05) clusters within the data. TFCE applies a weighted average between the cluster extent (i.e., number of connected samples) and cluster height (indicative of the magnitude of the statistical effect). This approach enables the detection of broad yet comparatively subtle clusters, which may be missed in conventional cluster permutation tests. Further, TFCE values were adjusted based on the membership status of individual data points within identified clusters, ensuring a nuanced scaling that accounts for the spatial extent of clusters.

### Code accessibility

The custom MATLAB scripts used in the current study are available to view on the Open Science Framework: https://osf.io/4yu9n/?view_only=68bd8912dcb2497ab688f33ebc6c13d3.

## Results

### Behavioural data

On Go trials, both the dominant (831.0 ± 14.6 ms) and non-dominant (827.0 ± 15.3 ms) hands lifted slightly after the target line on average. On Stop trials, stopping success rates of 51.2 ± 2.4 % indicate that the SSD staircasing procedure was effective. Mean SSRT was 210.5 ± 13.9 ms, reflecting response inhibition performance. Behavioural performance on Go and Stop trials was therefore as expected for healthy older adults and in keeping with previous studies using the ARIT (Hall et al., 2022; Macdonald et al., 2012).

### β-features

#### β-power

The LMM revealed a relative increase in average β-power primarily over central and right frontal electrodes during successful Stop trials compared to on-target Go trials (*p* < .05, TFCE-corrected; Fig. 2). At an individual electrode level, this significant finding reflected reduced β-desynchronisation in successful Stop trials, in contrast to larger β-desynchronisation in Go trials as participants prepared to move (Fig. 3A). This relative right frontal enhancement of β-power began to emerge around 188–156 ms prior to the SSRT, aligning with engagement of the right-lateralised inhibitory control network during successful stopping.

**Figure 3.**
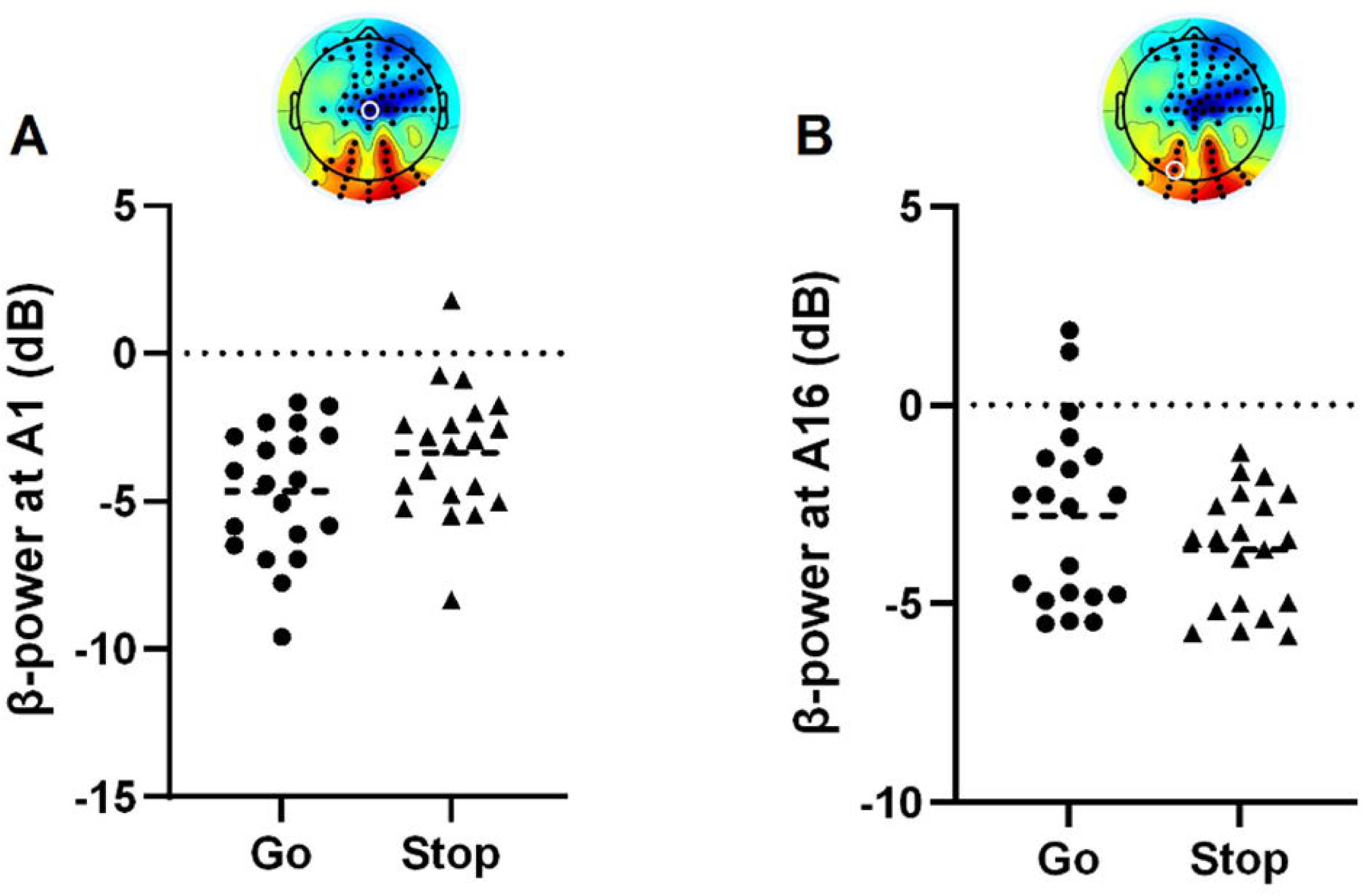
Individual-electrode β-power before response initiation and inhibition. In the final time bin (31 ms – lift-time/SSRT), panel A shows β-power at a central electrode (A1), demonstrating reduced β-desynchronisation in successful Stops compared to Go trials. Conversely, Panel B demonstrates relatively increased β-desynchronisation over a posterior electrode (A27) as participants inhibit the response in Stop trials.

In contrast, a relative decrease in average β-power over posterior (parieto-occipital) electrodes was established during successful Stop trials relative to on-target Go trials (*p* < .05). This activity originated in the right parietal region around 93–62 ms before the SSRT and subsequently spread to bilateral parietal and occipital areas, perhaps reflecting suppression of background sensory noise to focus inhibitory processes in Stop trials.

#### β-bursting

In support of our hypothesis, inhibition of the prepared response in Stop trials was characterised by a relative increase in fronto-central β-bursting (Fig. 4A); whereas β-bursting decreased over posterior areas (Fig. 4B). Interestingly, β-burst dynamics showed distinct spatial and temporal patterns to average β-power. First, while β-bursting also increased at right-lateralised frontal sites around 63–31 ms before the SSRT, it subsequently focalised to a more central motor hotspot (in line with our hypothesis) with activity extending to bilateral frontal electrodes as the SSRT approached. In contrast, increases in average β-power remained diffusely distributed across right frontal electrodes. Second, during successful response inhibition, early-occurring β-bursting was apparent at bilateral parieto-occipital electrodes (125–62 ms prior to SSRT; *p* < .05, TFCE-corrected; Fig. 2), suggesting increased transient β-activity in regions associated with attentional engagement. This pattern was not reflected in average β-power, which instead demonstrated reduced activation of these sites as the SSRT approached in Stop trials. Finally, there were also differences in β-burst activity relating to early time bins. During the response withholding phase, there was a relative increase in right fronto-central and sensorimotor β-bursting in Go trials (which was absent in average β-power) which subsequently decreased, perhaps reflecting early preparatory suppression processes.

**Figure 4.**
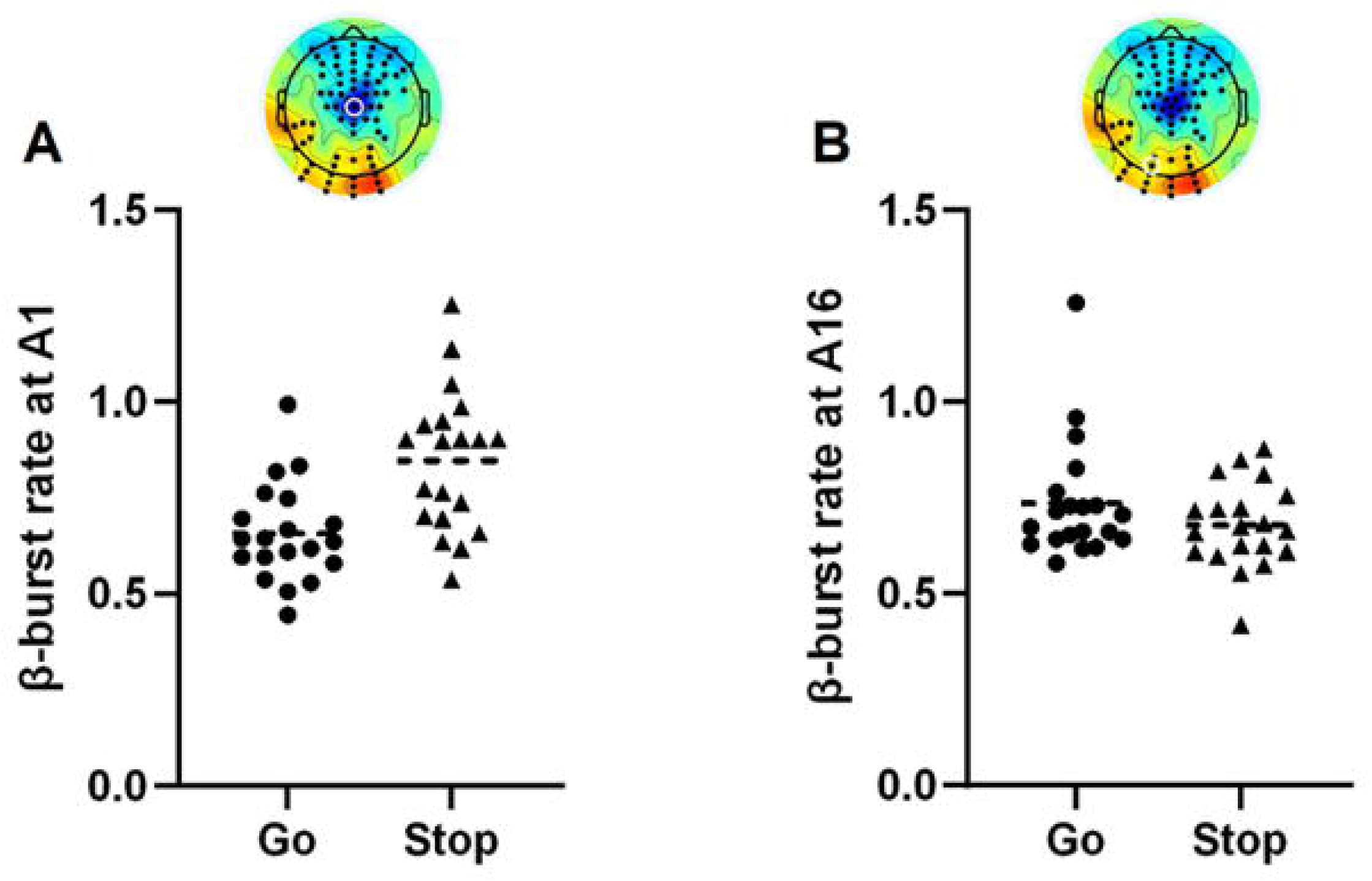
Individual-electrode β-burst rate before response initiation and inhibition. In the final time bin (31 ms – lift-time/SSRT), panel A shows β-burst rate at a central electrode (A1), demonstrating relatively increased β-burst rate in successful Stops compared to Go trials. Conversely, panel B shows a relative decrease in β-burst rate at a posterior electrode (A27) as participants inhibit the response in Stop trials.

β-burst duration mirrored the observed β-burst rate patterns (Fig. 2), with longer burst durations observed at (right) frontal and central electrodes in later time windows during successful Stop trials (further supporting our hypothesis) and early time windows during Go trials (*p* < .05, TFCE-corrected), reinforcing their role in inhibitory control and preparatory suppression, respectively. In early timepoints, the relative increase in right fronto-central and sensorimotor β-bursting in Go trials was more pronounced in β-burst duration than rate, perhaps indicating a stronger link between burst duration and preparatory suppression mechanisms. However, later in the trial, β-burst duration showed less pronounced parietal activation during Go trials and therefore may be somewhat less sensitive to attentional processes compared to burst rate.

### Muscle bursting

Following data collection, three participants were excluded from the muscle bursting analyses due to low signal-to-noise ratio in the EMG. As expected, healthy older adults demonstrated both premature bursts prior to the lift response on Go trials (Fig. 5A) and partial bursts when the lift would have occurred on successful Stop trials (Fig. 5B).

**Figure 5.**
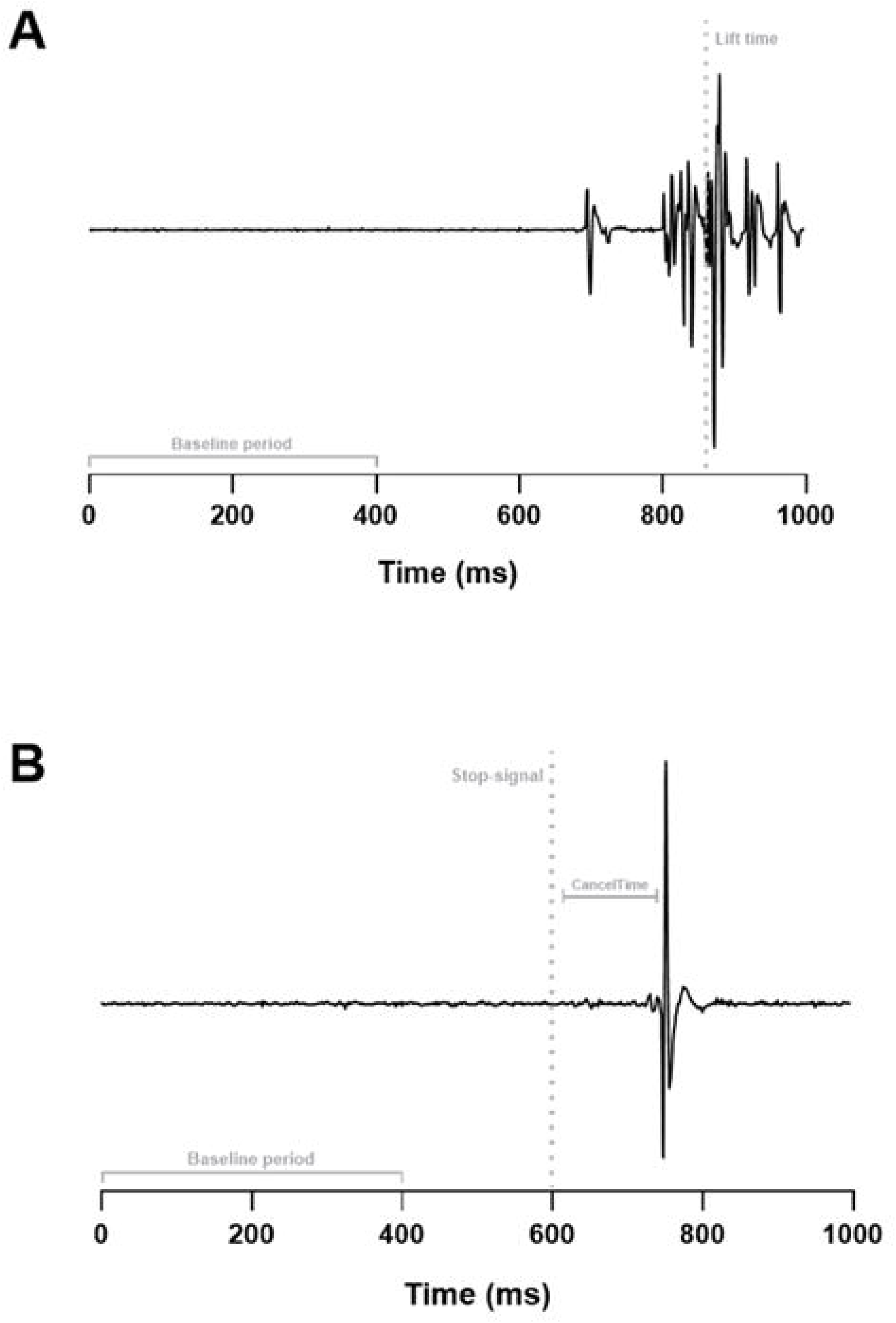
Representative EMG traces of muscle bursting. Panel A demonstrates a premature muscle burst (∼700 ms) prior to the main lift response (dotted line indicates lift-time). Panel B demonstrates a partial muscle burst following the stop-signal (dotted line) around when the response would have occurred (∼750-800 ms).

During response withholding, 18.4 ± 10.0% of Go trials presented with at least one premature burst, occurring 67–181 ms before the target on average (i.e., ∼100–210 ms before the lift-time). During response inhibition, 51.5 ± 20.4% of successful Stop trials presented with at least one partial burst. Mean CancelTime was 135.2 ± 16.6 ms (∼75 ms before SSRT). As expected, there was a positive correlation between CancelTime and SSRT (*r =* .578, *p* < .001).

### Single-trial integration of β- and muscle bursts

No significant clusters were identified for β-burst rate or duration between on-target Go trials which did or did not present premature muscle bursting. In contrast, the LMM revealed a significant increase in β-burst rate (but not duration) in a small cluster of electrodes (B17-18, B23) over right motor areas in successful Stop trials with partial muscle bursts compared to those without (*p* < .05, TFCE-corrected; Fig. 6). This effect was specific to the final time bin (31 ms – SSRT), occurring in close temporal proximity to inhibition of the anticipated response and supporting a functional link between β- and muscle bursting activity.

**Figure 6.**
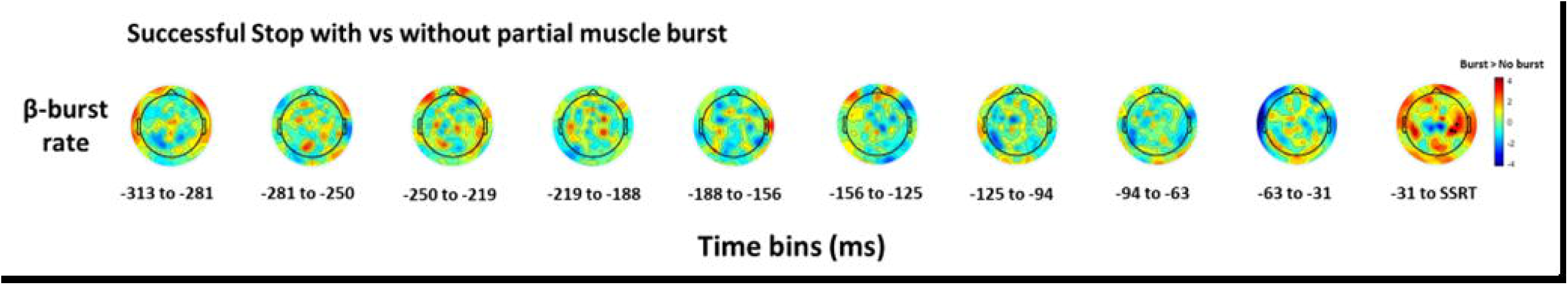
Topographical comparison of β-burst rate in Stop trials with and without a partial muscle burst. Topoplots of T-statistics for β-burst rate across ten ∼31 ms time bins leading up to the stop-signal reaction time (SSRT) on successful Stop trials with versus those without a partial muscle burst. Black dots refer to significant electrodes following threshold-free cluster enhancement permutation testing (p < .05). As per the colour-bar, red indicates a relative increase in β-burst rate in trials with a partial burst over the final time bin.

### Post-hoc temporal analysis

Following the observed link between β-burst rate and partial muscle bursting during successful Stop trials, we conducted a post-hoc temporal analysis to explore the relative timing of β- and muscle bursting. Using custom MATLAB scripts (2024b, The MathWorks), the time bin in which each partial muscle burst occurred was extracted, alongside the directly adjacent time bins (directly before and after). Trials with multiple partial bursts and those with partial bursts occurring in the final time bin (31 ms – SSRT) were therefore excluded from the analysis.

One-way ANOVAs were performed on β-burst rate values obtained in significant clusters identified in previous LMMs. There was no main effect of Time (*F*_2,48_ = 1.4, *p* = .249) on β-burst rate over the significant cluster of right motor electrodes linked with partial muscle bursting in the final time bin (Fig. 6). There was main effect of Time (*F*_2,48_ = 4.3, *p* = .019, □^2^ = .152) over the central β-bursting hotspot observed during response inhibition (Fig. 7), which demonstrated significantly increased β-burst rate in the time bin following partial muscle bursts (0.28 ± 0.08), compared to the time bin in which the burst occurred (0.23 ± 0.04; *p* = .04) and the prior time bin (0.23 ± 0.04; *p* = .047). β-burst rate was comparable in the time bins before and during muscle burst activity (*p* = 1.00).

**Figure 7.**
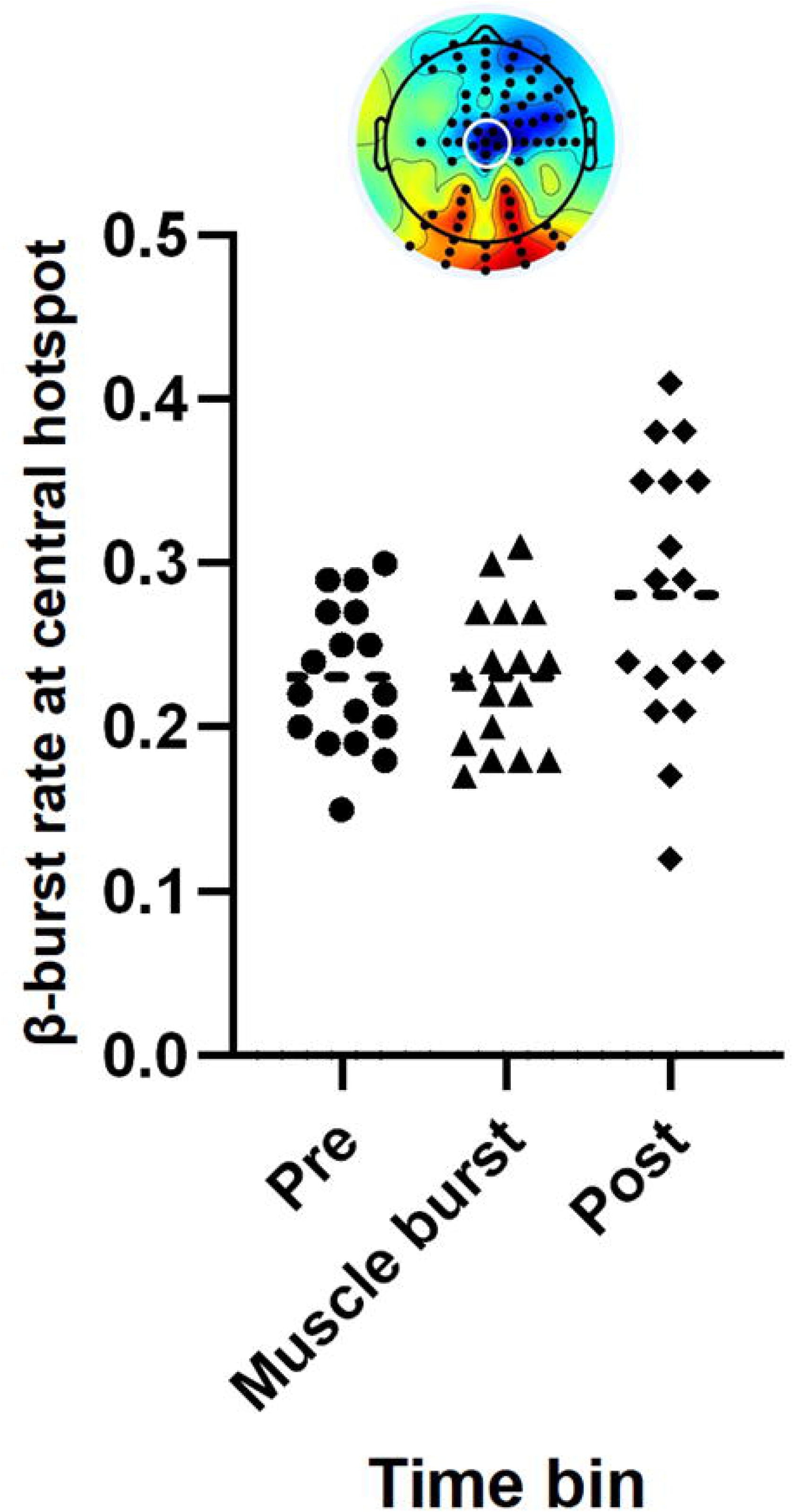
Mean β-burst rate over a central hotspot relative to muscle bursting. On successful Stop trials, a relative increase in β-bursting over central electrodes (A1-2, B1, C1, D1, D15) was apparent in the time bin following a partial burst, compared to before and during muscle bursting.

## Discussion

The present study applied an LMM-TFCE framework to map β-burst activity during the ARIT and relate these neural events to muscle bursts in healthy older adults. Our approach clearly elucidated distinct β-bursting patterns during response inhibition and withholding and uncovered a functional link between cortical β-bursting and late-stage gating of corticomotor drive. During response inhibition, we revealed left prefrontal activation in addition to the expected right fronto-central and bilateral sensorimotor activity characteristic of successful inhibitory control. This finding suggests our β-burst analysis was sensitive to broader inhibitory network recruitment during bimanual response cancellation.

Our data-driven approach examining brain-wide β-bursting further identified transient increases in bilateral parieto-occipital activity, likely related to stop-signal processing before a functional shift from attentional to inhibitory networks. Our β-bursting measures also proved uniquely sensitive to early fronto-central and sensorimotor activity during Go trials, perhaps reflecting tonic suppression preventing premature movement during response withholding.

Collectively, we provide evidence that β-bursting is a temporally precise neural marker of inhibitory control processes in healthy ageing. Combining β-burst detection with LMM-TFCE statistics is a viable analysis framework to examine sequences of transient oscillatory events and extend β-bursting research into heterogeneous populations. Our β-burst measures therefore hold potential as sensitive markers of age-related and clinical changes to inhibitory function.

### Response inhibition

Converging evidence supports a link between right frontal β-activity during response inhibition, as observed in the current study, and the right inferior frontal cortex (rIFC) (Swann et al., 2009; Swann et al., 2012). Disruption of the rIFC around β-burst onset prolongs stopping latency (Hannah et al., 2020; Sundby et al., 2021), positioning the rIFC as a ‘brake’ for motor output triggered by an unexpected event (e.g., the stop-signal) (Aron et al., 2014). The rIFC may initiate a cascade of neural processes transmitted via inhibitory β-oscillations through the pre-supplementary motor area (pre-SMA) (Schaum et al., 2021), culminating in pre-motor inhibition of M1 to suppress motor output (Fiori et al., 2016). This reinforces the role of right frontal β-band signalling driven by the rIFC as a neural signature of inhibitory control.

While average β-power demonstrated stereotypical right fronto-central activity during response inhibition, β-bursting uncovered bilateral frontal activation. Often absent in response inhibition literature, select studies have linked the left IFC (lIFC) and pre-SMA to bimanual inhibition (Fiori et al., 2016; Schaffer et al., 2020; Swick et al., 2008). The robust analysis in the present study may have facilitated the identification of bilateral frontal contributions, implicating integrated contralateral and ipsilateral pathways involving the lIFC and pre-SMA in the inhibition of an internally generated bimanual response (Zhang et al., 2017). However, bilateral frontal activity may also be interpreted as age-related compensatory mechanisms, where increased lIFG engagement may support declining right-lateralised prefrontal function (Berlingeri et al., 2013; Cabeza, 2002). Either way, these findings position β-bursting detected by our analysis framework as a sensitive and mechanistically informative measure of inhibitory control, revealing task- and perhaps age-dependent neural dynamics.

Aside from frontal activity, the observed temporal fluctuations in posterior β-bursting during response inhibition could be linked to attentional networks. An initial increase in parieto-occipital β-bursting was observed following the stop-signal. Parietal activity in motor control tasks links with attentive processing of task-relevant stimuli (Boehler et al., 2010), indicative of selective attending to the unexpected stop-signal. However, as the SSRT approached, β-burst activity over these areas was suppressed. This aligns with work by Enz et al. (2020), reporting a relative decrease in β-burst volume in centroparietal and occipital areas during successful Stop trials of the Stop-Signal Task. The attenuation of posterior β-activity could minimise sensory or attentional ‘noise’, shifting toward optimised top-down inhibitory control.

In line with these temporally dynamic patterns, right sensorimotor β-bursting was linked with partial muscle bursting on successful Stop trials, suggestive of a fast-acting mechanism in response to activation of the go process. Temporal analyses further supported this theory, demonstrating increased central β-bursting following a partial burst. We speculate that cortical inhibitory mechanisms, such as dynamic response threshold modulation by the right pre-SMA (Wolpe et al., 2022), are exerted to rapidly suppress already-initiated motor activity. However, this analysis was exploratory. Trials with muscle bursts in the final time bin were excluded due to the lack of a subsequent bin for comparison. Temporal variability was also introduced, as muscle bursts occurred at varying points within each ∼32 ms time bin. While increased β-activity after muscle bursting could reflect general engagement of inhibitory control over time, the comparable activity between the prior two time bins suggests a functional link to the muscle burst itself. Future research could extend the post-trial epoch to capture β-burst dynamics following suppression at the muscle.

### Response withholding

Fronto-central and sensorimotor β-bursting was initially elevated during response withholding in Go trials. This may reflect a tonically inhibited motor state, indexing preparatory suppression mechanisms from the IFC/pre-SMA, via the basal ganglia to intracortical M1 networks (Aron & Poldrack, 2006; Coxon et al., 2006) to prevent premature response initiation. Such mechanisms could not be discerned from average β-power. Subsequently, as the lift response was initiated, fronto-central and sensorimotor β-activity reduced, which aligns with a preparatory release of intracortical inhibition in M1 to facilitate a rapid response (Davranche et al., 2007; Hannah et al., 2018; Sinclair & Hammond, 2008, 2009). It should be acknowledged that comparable response withholding is expected before stop-signal presentation in Stop trials. The observed differences between trial types could be due to time-locking β-bursts to the end of the individualised going/stopping process. While allowing direct comparison of response initiation and inhibition, this approach compromised temporal precision earlier in Stop trials, where the stop-signal was presented at variable times. It may therefore have been difficult to isolate the neural dynamics underlying response withholding during Stop trials.

There was no evidence of a relationship between β-bursting activity and premature muscle bursts during Go trials. It is possible that modulation of inhibitory control is not responsible for these ineffective muscle bursts during response withholding. Alternatively, two methodological constraints may have masked this relationship. Firstly, there was a low rate of premature bursts on Go trials (4–39%). The LMM analyses integrating β-bursting and muscle bursting were contingent on the presence of identifiable muscle bursts. Secondly, current β-burst detection methods are optimised for fronto-central activity associated with reactive inhibition. Future research should more directly examine the mechanisms of withholding an internally generated response to better capture its distinct timing and neural characteristics.

### Clinical implications

The current findings support combining β-burst detection with LMM-TFCE statistics to examine β-bursting patterns in heterogeneous populations. This approach could augment β-bursting analyses across a range of research contexts, including neural development (Rayson et al., 2023). More specifically, our β- (and muscle) bursting measures have potential as trial-by-trial markers of inhibitory function for characterising impulse control issues in clinical populations. Individuals with Parkinson’s disease exhibit elevated rates of premature and partial muscle bursts, alongside maladaptive modulation of corticomotor excitability, which is associated with impaired ARIT performance (Warden et al., 2025). These neurophysiological changes may reflect disrupted top-down β-burst signalling, compromising GABAergic M1 circuits, thereby failing to sufficiently suppress corticomotor excitability and resultant motor activity. Future research could investigate whether the observed β-burst patterns linked with response withholding and inhibition differ in clinical populations, evaluating their potential as objective markers of impulse control dysfunction.

### Potential considerations

Due to analysis constraints, we could not report β-burst amplitude and volume. These measures showed high standard error around the head, likely from reduced β-desynchronisation, which lead to a poor regression fit between T-statistics and β-coefficients. Increasing the burst threshold (2× to 6× median) did not resolve these inconsistencies, so amplitude and volume measures were excluded to preserve data integrity. However, β-burst volume may better predict successful inhibition than β-burst rate (Enz et al., 2021), highlighting its potential relevance to muscle bursting dynamics in future work.

We revealed a temporal relationship between β-bursting and partial muscle bursts; however, the β-burst analyses were time-locked to the SSRT. The SSRT provides an estimate of stopping performance based on average behavioural data. Employing CancelTime as the stopping outcome measure in future could yield a more precise trial-by-trial indicator of stopping latency, to better characterise the temporal dynamics of inhibitory control.

## Conclusion

The current study highlights β-bursting as a temporally precise neural marker of inhibitory control in healthy older adults. Our novel use of a robust LMM-TFCE analysis framework identified distinct β-bursting patterns associated with both proactive response withholding and reactive inhibition, which were not captured by averaged β-power. These results underscore the sensitivity of the applied β-bursting analysis to both the timing and context of inhibitory demands, supporting its utility as a mechanistic and clinically relevant marker of inhibitory function.

## Data availability

The anonymised datasets analysed in the current study are available from the corresponding author upon reasonable request. Access will be granted in accordance with institutional guidelines and subject to ethical approval where required.

## Acknowledgements

This work was supported by the Les Rhoades Memorial PhD Studentship from the Humane Research Trust. The authors gratefully acknowledge the commitment of participants who contributed their time to this study. The authors would also like to acknowledge Birmingham 1000 Elders for their support with recruitment, and Brogan Ling and Joshua Pearson for assistance with data collection.

During the preparation of this work the authors used Microsoft Copilot to make some writing more succinct. After using this tool/service, the authors reviewed and edited the content as needed and take full responsibility for the content of the publication.

## References

Aron, A. R., Herz, D. M., Brown, P., Forstmann, B. U., & Zaghloul, K. (2016). Frontosubthalamic Circuits for Control of Action and Cognition. J Neurosci, 36(45), 11489–11495. 10.1523/jneurosci.2348-16.2016

Aron, A. R., & Poldrack, R. A. (2006). Cortical and subcortical contributions to Stop signal response inhibition: role of the subthalamic nucleus. J Neurosci, 26(9), 2424–2433. 10.1523/jneurosci.4682-05.2006

Aron, A. R., Robbins, T. W., & Poldrack, R. A. (2014). Inhibition and the right inferior frontal cortex: one decade on. Trends Cogn Sci, 18(4), 177–185. 10.1016/j.tics.2013.12.003

Berlingeri, M., Danelli, L., Bottini, G., Sberna, M., & Paulesu, E. (2013). Reassessing the HAROLD model: is the hemispheric asymmetry reduction in older adults a special case of compensatory-related utilisation of neural circuits? Exp Brain Res, 224(3), 393–410. 10.1007/s00221-012-3319-x

Boehler, C. N., Appelbaum, L. G., Krebs, R. M., Hopf, J. M., & Woldorff, M. G. (2010). Pinning down response inhibition in the brain — Conjunction analyses of the Stop-signal task. NeuroImage, 52(4), 1621–1632. 10.1016/j.neuroimage.2010.04.276

Cabeza, R. (2002). Hemispheric asymmetry reduction in older adults: the HAROLD model. Psychol Aging, 17(1), 85–100. 10.1037//0882-7974.17.1.85

Coxon, J. P., Stinear, C. M., & Byblow, W. D. (2006). Intracortical inhibition during volitional inhibition of prepared action. J Neurophysiol, 95(6), 3371–3383. 10.1152/jn.01334.2005

Davranche, K., Tandonnet, C., Burle, B., Meynier, C., Vidal, F., & Hasbroucq, T. (2007). The dual nature of time preparation: neural activation and suppression revealed by transcranial magnetic stimulation of the motor cortex. European Journal of Neuroscience, 25(12), 3766–3774.

Delorme, A., & Makeig, S. (2004). EEGLAB: an open source toolbox for analysis of single-trial EEG dynamics including independent component analysis. J Neurosci Methods, 134(1), 9–21. 10.1016/j.jneumeth.2003.10.009

Enz, N., Ruddy, K. L., Rueda-Delgado, L. M., & Whelan, R. (2021). Volume of β-Bursts, But Not Their Rate, Predicts Successful Response Inhibition. J Neurosci, 41(23), 5069–5079. 10.1523/jneurosci.2231-20.2021

Fiori, F., Chiappini, E., Soriano, M., Paracampo, R., Romei, V., Borgomaneri, S., & Avenanti, A. (2016). Long-latency modulation of motor cortex excitability by ipsilateral posterior inferior frontal gyrus and pre-supplementary motor area. Scientific Reports, 6(1), 38396. 10.1038/srep38396

Groth, C. L., Singh, A., Zhang, Q., Berman, B. D., & Narayanan, N. S. (2021). GABAergic Modulation in Movement Related Oscillatory Activity: A Review of the Effect Pharmacologically and with Aging. Tremor Other Hyperkinet Mov (N Y), 11, 48. 10.5334/tohm.655

Hall, A., Jenkinson, N., & MacDonald, H. J. (2022). Exploring stop signal reaction time over two sessions of the anticipatory response inhibition task. Exp Brain Res, 240(11), 3061–3072. 10.1007/s00221-022-06480-x

Hannah, R., Cavanagh, S. E., Tremblay, S., Simeoni, S., & Rothwell, J. C. (2018). Selective Suppression of Local Interneuron Circuits in Human Motor Cortex Contributes to Movement Preparation. J Neurosci, 38(5), 1264–1276. 10.1523/jneurosci.2869-17.2017

Hannah, R., Muralidharan, V., Sundby, K. K., & Aron, A. R. (2020). Temporally-precise disruption of prefrontal cortex informed by the timing of beta bursts impairs human action-stopping. NeuroImage, 222, 117222. 10.1016/j.neuroimage.2020.117222

He, J. L., Hirst, R. J., Puri, R., Coxon, J., Byblow, W., Hinder, M., Skippen, P., Matzke, D., Heathcote, A., Wadsley, C. G., Silk, T., Hyde, C., Parmar, D., Pedapati, E., Gilbert, D. L., Huddleston, D. A., Mostofsky, S., Leunissen, I., MacDonald, H. J., . . . Puts, N. A. J. (2022). OSARI, an Open-Source Anticipated Response Inhibition Task. Behav Res Methods, 54(3), 1530–1540. 10.3758/s13428-021-01680-9

Hodges, P. W., & Bui, B. H. (1996). A comparison of computer-based methods for the determination of onset of muscle contraction using electromyography. Electroencephalogr Clin Neurophysiol, 101(6), 511–519. 10.1016/s0013-4694(96)95190-5

Jana, S., Hannah, R., Muralidharan, V., & Aron, A. R. (2020). Temporal cascade of frontal, motor and muscle processes underlying human action-stopping. Elife, 9, e50371. 10.7554/eLife.50371

Karlawish, J., Cary, M., Moelter, S. T., Siderowf, A., Sullo, E., Xie, S., & Weintraub, D. (2013). Cognitive impairment and PD patients’ capacity to consent to research. Neurology, 81(9), 801–807. 10.1212/WNL.0b013e3182a05ba5

Kayser, J., & Tenke, C. E. (2006). Principal components analysis of Laplacian waveforms as a generic method for identifying ERP generator patterns: I. Evaluation with auditory oddball tasks. Clin Neurophysiol, 117(2), 348–368. 10.1016/j.clinph.2005.08.034

Kilavik, B. E., Zaepffel, M., Brovelli, A., MacKay, W. A., & Riehle, A. (2013). The ups and downs of beta oscillations in sensorimotor cortex. Experimental Neurology, 245, 15–26. 10.1016/j.expneurol.2012.09.014

Little, S., Bonaiuto, J., Barnes, G., & Bestmann, S. (2019). Human motor cortical beta bursts relate to movement planning and response errors. PLoS Biol, 17(10), e3000479. 10.1371/journal.pbio.3000479

MacDonald, H. J., Coxon, J. P., Stinear, C. M., & Byblow, W. D. (2014). The fall and rise of corticomotor excitability with cancellation and reinitiation of prepared action. J Neurophysiol, 112(11), 2707–2717. 10.1152/jn.00366.2014

Macdonald, H. J., Stinear, C. M., & Byblow, W. D. (2012). Uncoupling response inhibition. J Neurophysiol, 108(5), 1492–1500. 10.1152/jn.01184.2011

Mensen, A., & Khatami, R. (2013). Advanced EEG analysis using threshold-free cluster-enhancement and non-parametric statistics. NeuroImage, 67, 111–118. 10.1016/j.neuroimage.2012.10.027

Meyer, H. C., & Bucci, D. J. (2016). Neural and behavioral mechanisms of proactive and reactive inhibition. Learn Mem, 23(10), 504–514. 10.1101/lm.040501.115

Mognon, A., Jovicich, J., Bruzzone, L., & Buiatti, M. (2011). ADJUST: An automatic EEG artifact detector based on the joint use of spatial and temporal features. Psychophysiology, 48(2), 229–240. 10.1111/j.1469-8986.2010.01061.x

Nolan, H., Whelan, R., & Reilly, R. B. (2010). FASTER: Fully Automated Statistical Thresholding for EEG artifact Rejection. J Neurosci Methods, 192(1), 152–162. 10.1016/j.jneumeth.2010.07.015

Raud, L., Thunberg, C., & Huster, R. J. (2022). Partial response electromyography as a marker of action stopping. Elife, 11. 10.7554/eLife.70332

Rayson, H., Szul, M. J., El-Khoueiry, P., Debnath, R., Gautier-Martins, M., Ferrari, P. F., Fox, N., & Bonaiuto, J. J. (2023). Bursting with potential: how sensorimotor beta bursts develop from infancy to adulthood. Journal of Neuroscience, 43(49), 8487–8503.

Rey-Mermet, A., & Gade, M. (2018). Inhibition in aging: What is preserved? What declines? A meta-analysis. Psychon Bull Rev, 25(5), 1695–1716. 10.3758/s13423-017-1384-7

Riccardi, N., Teghipco, A., Newman-Norlund, S., Newman-Norlund, R., Rangus, I., Rorden, C., Fridriksson, J., & Bonilha, L. (2025). Distinct brain age gradients across the adult lifespan reflect diverse neurobiological hierarchies. Communications Biology, 8(1), 802. 10.1038/s42003-025-08228-z

Schaffer, J. E., Maenza, C., Good, D. C., Przybyla, A., & Sainburg, R. L. (2020). Left hemisphere damage produces deficits in predictive control of bilateral coordination. Exp Brain Res, 238(12), 2733–2744. 10.1007/s00221-020-05928-2

Schaum, M., Pinzuti, E., Sebastian, A., Lieb, K., Fries, P., Mobascher, A., Jung, P., Wibral, M., & Tüscher, O. (2021). Right inferior frontal gyrus implements motor inhibitory control via beta-band oscillations in humans. Elife, 10. 10.7554/eLife.61679

Sherman, M. A., Lee, S., Law, R., Haegens, S., Thorn, C. A., Hämäläinen, M. S., Moore, C. I., & Jones, S. R. (2016). Neural mechanisms of transient neocortical beta rhythms: Converging evidence from humans, computational modeling, monkeys, and mice. Proceedings of the National Academy of Sciences, 113(33), E4885–E4894.

Sinclair, C., & Hammond, G. R. (2008). Reduced intracortical inhibition during the foreperiod of a warned reaction time task. Experimental Brain Research, 186, 385–392.

Sinclair, C., & Hammond, G. R. (2009). Excitatory and inhibitory processes in primary motor cortex during the foreperiod of a warned reaction time task are unrelated to response expectancy. Experimental Brain Research, 194, 103–113.

Sundby, K. K., Jana, S., & Aron, A. R. (2021). Double-blind disruption of right inferior frontal cortex with TMS reduces right frontal beta power for action stopping. Journal of Neurophysiology, 125(1), 140–153.

Swann, N., Tandon, N., Canolty, R., Ellmore, T. M., McEvoy, L. K., Dreyer, S., DiSano, M., & Aron, A. R. (2009). Intracranial EEG reveals a time-and frequency-specific role for the right inferior frontal gyrus and primary motor cortex in stopping initiated responses. Journal of Neuroscience, 29(40), 12675–12685.

Swann, N. C., Cai, W., Conner, C. R., Pieters, T. A., Claffey, M. P., George, J. S., Aron, A. R., & Tandon, N. (2012). Roles for the pre-supplementary motor area and the right inferior frontal gyrus in stopping action: electrophysiological responses and functional and structural connectivity. NeuroImage, 59(3), 2860–2870. 10.1016/j.neuroimage.2011.09.049

Swick, D., Ashley, V., Turken, & U. (2008). Left inferior frontal gyrus is critical for response inhibition. BMC Neuroscience, 9(1), 102. 10.1186/1471-2202-9-102

Verbruggen, F., Aron, A. R., Band, G. P., Beste, C., Bissett, P. G., Brockett, A. T., Brown, J. W., Chamberlain, S. R., Chambers, C. D., Colonius, H., Colzato, L. S., Corneil, B. D., Coxon, J. P., Dupuis, A., Eagle, D. M., Garavan, H., Greenhouse, I., Heathcote, A., Huster, R. J., . . . Boehler, C. N. (2019). A consensus guide to capturing the ability to inhibit actions and impulsive behaviors in the stop-signal task. Elife, 8. 10.7554/eLife.46323

Visalli, A., Montefinese, M., Viviani, G., Finos, L., Vallesi, A., & Ambrosini, E. (2024). lmeEEG: Mass linear mixed-effects modeling of EEG data with crossed random effects. Journal of Neuroscience Methods, 401, 109991. 10.1016/j.jneumeth.2023.109991

Warden, A. C. M., McAllister, C., Cruse, D., Wright, B., & MacDonald, H. J. (2025). Muscle bursting and corticomotor excitability index impaired impulse control in Parkinson’s disease. medRxiv, 2025.2004.2028.25326550. 10.1101/2025.04.28.25326550

Wessel, J. R. (2020). β-Bursts Reveal the Trial-to-Trial Dynamics of Movement Initiation and Cancellation. J Neurosci, 40(2), 411–423. 10.1523/jneurosci.1887-19.2019

Wolpe, N., Hezemans, F. H., Rae, C. L., Zhang, J., & Rowe, J. B. (2022). The pre-supplementary motor area achieves inhibitory control by modulating response thresholds. Cortex, 152, 98–108. 10.1016/j.cortex.2022.03.018

Zhang, R., Geng, X., & Lee, T. M. C. (2017). Large-scale functional neural network correlates of response inhibition: an fMRI meta-analysis. Brain Struct Funct, 222(9), 3973–3990. 10.1007/s00429-017-1443-x

